# Metabolic and transcriptomic insights into a GalNAc/Man-Specific lectin in yeast fermentation

**DOI:** 10.1101/2025.05.26.656115

**Authors:** Shuai Liu, Linge Li, Hongxin Niu, Wei Li, Changqing Tong

## Abstract

Brewer’s yeast (*Saccharomyces cerevisiae*) plays a central role in fermentation, and improving its performance is crucial in the brewing industry. This study explored how *Cyclina sinensis* lectin (CSL), a bioactive peptide from shellfish, influences yeast metabolism using proteomics and metabolomics. CSL bound to yeast peptidoglycan in a concentration-dependent manner and altered cell surface structure. Proteomic analysis revealed 117 differentially expressed proteins after 24 h of CSL treatment, with upregulation of key glycolytic enzymes (e.g., hexokinase, GAPDH) and downregulation of TCA cycle enzymes, suggesting enhanced ethanol production via glycolysis activation and TCA suppression. Metabolomic profiling further confirmed this, showing increased α-D-glucose, glucose-6-phosphate, NAD^+^, and glutathione, alongside reduced TCA intermediates like malic acid. These results indicate CSL promotes ethanol accumulation by reprogramming central carbon metabolism. This study provides novel insight into the metabolic modulation of yeast by marine lectins and suggests potential applications of CSL in improving fermentation

## Introduction

Recent years, lectins have become an important research area of interest in disciplines such as glycobiology, immunology and microbiology, because of their wide range of bioactivities. Thousands of lectins have been discovered so far, and it is noteworthy to discover lectins with special physiological activities that have novel structures. Marine invertebrates, such as C*yclina sinensis*, often produce metabolites with special structures due to the special living environment, including novel lectins.

C*yclina sinensis*, commonly known as the Chinese venus, is a species of bivalve mollusk in the family Veneridae which is mainly distributed in China, Japan and Korea (*1*). Currently, there are not many studies on the active substances of it. *Cyclina sinensis* have high nutritional value, rich in protein, fat, carbohydrates, sugar, inorganic elements, etc (*2*). *Cyclina sinensis* contain a large number of polysaccharide active substances. Lectins are a class of glycoproteins with multivalent sugar-binding properties and have diverse physiological activities. A N-acetyl-galactosamine/mannose-specific lectin (CSL) with a molecular weight of 72 kDa has been isolated from the haemolymph of the *Cyclina sinensis* (*3*). CSL has the ability to promote immune response, exhibit antioxidant, antitumor, and antibacterial properties, stimulate yeast cell proliferation and growth, agglutinate yeast, and enhance ethanol production (*3–7*). Industrial extraction process of *Cyclina sinensis* polysaccharide is mature and its biological activity is clear and suitable for mass production preparation, which is a potential source of health food and pharmaceuticals.

Yeast is the collective name of a group of unicellular fungi, which is one of the earliest and most important microbial strains used in the practice of human production and life. Among them, *Saccharomyces cerevisiae* is the most typical yeast species, and is also a commonly used microorganism for fermentation and baking in the food industry. *Saccharomyces cerevisiae* produces ethanol by fermenting glucose under anaerobic conditions, and is therefore widely used in the brewing of beer, liquor, fruit wines and the manufacture of bread (*8,9*). *Saccharomyces cerevisiae* is a unicellular parthenogenetic anaerobic microorganism belonging to the class of fungi, which has a nucleus, a cell membrane, a cell wall and mitochondria. During the growth of *Saccharomyces cerevisiae*, at least 20 different proteins are present in the outer protein layer of the cell wall (*10–12*). Thus, the yeast cell wall has the basis for producing a wide variety of physiological functions. Cell wall proteins play a crucial role in yeast cell flocculation, cross type, membrane formation, mycelium formation, and invasive growth (*10,13,14*) . The structure of the cell wall of *Saccharomyces cerevisiae* is relatively stable. Thus, the yeast cell wall has the basis for producing a wide variety of physiological functions. Yeast fermentation is affected by many factors, such as temperature, agitation rate, medium composition, precursor amino acid addition, glucose streaming and addition strategy, pH of the fermentation substrate and exogenous additives. Research has been carried out on the modification of yeast fermentation performance through the addition of exogenous substances. The selection of brewer’s yeast and the improvement of brewer’s technology are two important factors to improve ethanol production by brewer’s yeast fermentation. For the improvement of brewing technology, i.e., to enhance the proliferation and fermentation performance of brewer’s yeast, the addition of peptide actives is a potential method with both theoretical and applied values, especially the addition of newly discovered lectin-like actives. Tong et al. found that Colocasia esculenta Lectin (CAL) isolated from sponge (*Craniella australiensis*) had a stimulatory effect on the fermentation of *Saccharomyces cerevisiae*, and the ethanol produced increased when the concentration of CAL increased (*15*). Jin et al. investigated the effect of N-acetylgalactosamine/galactose-specific lectin (CGL) isolated from mussels (*Crenomytilus grayanus*) promoted agglutinating and growth of yeast (*16*). All these lectins mentioned above had a promotional effect on the yeast fermentation process.

Lectins are non-enzymatic, non-immunogenic and multivalent sugar-binding proteins with diverse sugar recognition specificities. The proteins can form complexes with the yeast cell wall, a process that is the beginning of the protein’s influence on yeast physiological processes (*17*).

The binding of lectins and sugars on the surface of yeast cells drives the yeast to agglutination, which has a great influence on yeast fermentation. Yeast cells in the agglutinated state have better resistance to environmental stresses and have a longer lifespan (*18*). *Cyclina sinensis* lectin is a new type of sugar-binding protein with the ability to interact with yeast (*3*). *Cyclina sinensis* is a shellfish that is relatively large farmed, and its lectin is more readily available. By studying the interaction of its lectin with *Saccharomyces cerevisiae*, it is of great significance to understand the signal transduction pathway of lectin binding to *Saccharomyces cerevisiae* cells. Meanwhile, the results of the study also provide basic data for the potential application of *Cyclina sinensis* lectins in the fermentation process of yeast.

The rapid development of high-throughput protein separation and identification technology has brought great changes to proteomic technology. Mass spectrometry has become the core technology of proteomic technology (*19*). Proteomics studies the landscape of protein expression at a specific time. Aiming at the characterizing a large variety of small metabolites with a wide span of polarity and concentration dynamics, liquid chromatography-mass spectrometry (LC-MS) technology has become one of the most important tools (*20,21*) . With the combination of proteomic and metabolomic technologies, the mechanism of lectin regulation of proliferation and metabolism in *Saccharomyces cerevisiae* could be advanced. In this paper, we have studied the interactions between CSL and yeast cell walls and membranes using biochemical, proteomic and metabolomic techniques. We aimed to elucidate the protein-level regulation of *Saccharomyces cerevisiae* proliferation and metabolism and verify the effects of CSL on yeast fermentation using metabolomic techniques, providing a foundation for its application in fermentation.

## Materials & methods

### Sample collection

We obtained *Cyclina sinensis* from a local market. *Saccharomyces cerevisiae* was purchased from Angel Yeast Co., horse radish peroxidase (HRP) and bovine serum albumin (BSA) were obtained from Beijing Solarbio Science & Technology Co.,Ltd. Peptidoglycan from *S. cerevisiae*, mannose, and N-acetyl-D-galactosamine were sourced from Sigma, USA. The YPD liquid medium was prepared by dissolving 2 g of dextrose, 1 g of yeast extract, and 2 g of peptone in 100 mL of deionized water, followed by sterilization at 121°C for 20 minutes. Other analytical-grade reagents were purchased from commercial suppliers. The instruments used in this study included a Triple TOF 6600^+^ mass spectrometer (AB SCIEX), an Agilent 1290 Infinity LC system (Agilent), a UV-1750 spectrophotometer (Shimadzu), a pHS-3C pH meter (Shanghai INESA Scientific Instruments Co.), and a 5430R centrifuge (Eppendorf).

Fresh fringe tissues of razor clams were selected, ground into a slurry, and extracted with physiological saline (0.9 g NaCl per 100 mL of slurry). The extract was incubated for 12 hours, centrifuged at 9000 rpm for 15 minutes, and the supernatant was collected. Ammonium sulfate was added to 80% saturation, and after another 12 hours of incubation, the precipitate was collected, dialyzed, and lyophilized to obtain razor clam lectin (CSL).

The metabolic products of the co-fermentation of *Cyclina sinensis* lectin and *S. cerevisiae* for 24 hours, as well as the fermentation products of *S. cerevisiae* alone for 24 hours, were used for further analysis.

### CSL isolation and purification

We isolated CSL from *C. sinensis* haemolymph using Cellulose DE52 ion-exchange chromatography, Sephadex G-100 molecular sieve chromatography, and HPLC, following the method described by Tong et al. (2012).

### Yeast fermentation and interaction with CSL

We prepared the fermented liquid medium containing150 g of glucose, 5 g of yeast dip powder and 10 g of peptone, the ingredients were dissolved in 1000 mL of deionized water and sterilized for 20 min at 121°C. A 500 mL yeast suspension (OD_600_ = 0.387±0.004) was prepared and mixed with 30 mL of 2.69 mg/mL CSL solution. We incubated the mixture in a shaker at 150 rpm and 30°C for 24 h. After fermentation, samples were centrifuged at 8500 rpm for 10 min at 4°C.The yeast precipitate was collected for metabolomics analysis, and ethanol content was measured in the supernatant. The assay was performed three times in parallel, with no CSL added to the control group.

### Illumina RNA-seq

We carried out RNA-seq using the Illumina platform. RNA-seq was performed using the Illumina sequencing platform at Shanghai ZK-Life Science. The RNA-seq workflow involved total RNA sample detection, mRNA enrichment, double-stranded cDNA synthesis, end repair, A-tailing, adapter ligation, fragment selection, PCR amplification, library quality control, and Illumina sequencing. First, the RNA degradation level and potential contamination were assessed using 1% agarose gel electrophoresis. The purity (OD_260_/OD_280_ ratio), concentration, and absorbance peaks of the RNA were measured using a Nanodrop spectrophotometer. RNA concentration and integrity were further analyzed using the Agilent 2100 Bioanalyzer with the RNA Nano 6000 assay kit (Agilent Technologies, USA). Key assessment parameters included RIN value, 28 S/18 S ratio, baseline stability, and presence of the 5 S peak. Raw data was evaluated by quality check and filter for following analysis.

### Metabolomics sample preparation

We grounded the *C. sinensis* in liquid nitrogen. And then 200 μL ultrapure water and 800 μL methanol/acetonitrile (1:1, v/v) were added. Samples were vortexed, ultrasonicated at low temperature for 30 min, incubated at -20°C for 1 h, and centrifuged at 13000 rpm for 15 min at 4°C.The supernatant was lyophilized and stored at -80°C.

### Chromatography

We conducted Chromatographic analysis using a hydrophilic interaction liquid chromatography (HILIC) column maintained at 25°C. The mobile phase consisted of solvent A (water with 25 mmol/L ammonium acetate and 25 mmol/L ammonia) and solvent B (acetonitrile). The flow rate was set at 0.3 mL/min, and the injection volume was 2 μL. The gradient elution program was as follows: 85% B was maintained from 0 to 1 min, followed by a linear decrease from 85% to 65% B between 1 and 12 min. From 12 to 12.1 min, B further decreased linearly from 65% to 40%.

Then, B increased linearly from 40% to 85% between 12.1 and 15 min. At 15–15.1 min, B was maintained at 40%, and from 15.1 to 20 min, B was held at 85%. Throughout the analysis, samples were placed in a 4°C autosampler. Successive sample injections were performed in a randomized order, with quality control (QC) samples inserted at regular intervals to assess system stability and ensure data reliability.

### Mass spectrometry analysis

A quadrupole time-of-flight (Q-TOF) mass spectrometer equipped with an electrospray ionization (ESI) source was used for detection in both positive and negative ion modes. After UHPLC separation, mass spectrometric analysis was performed under the following conditions.

The electrospray ionization (ESI) source parameters were set as follows: ion source gas 1 (Gas1) and gas 2 (Gas2) were both maintained at 60, with a curtain gas (CUR) of 30. The source temperature was set at 600°C, and the ion spray voltage floating (ISVF) was ±5500 V for both positive and negative ion modes.

Mass spectrometric acquisition parameters were optimized as follows: the TOF MS scan range was m/z 60–1000 Da, and the product ion scan range was m/z 25–1000 Da. The accumulation time was set to 0.20 s per spectrum for TOF MS scans and 0.05 s per spectrum for product ion scans.

For MS/MS analysis, information-dependent acquisition (IDA) mode with high sensitivity was applied. The declustering potential (DP) was set at ±60 V for both positive and negative ion modes, with a collision energy (CE) of 35 ± 15 eV. Isotope exclusion was set within 4 Da, and up to six candidate ions were monitored per cycle.

### RNA-seq differential expression and functional enrichment analysis

For our study, we employed a differential expression (DE) analysis approach utilizing the edgeR (*22*) package to distinguish the genes that exhibited significant changes in expression across various conditions. The initial step involved transforming the raw read counts into counts per million (CPM), ensuring that only genes with a CPM value exceeding 0.5 in at least one sample across all four biological replicates were considered for further scrutiny. These CPM values underwent quantile normalization through the voom (*23*) function to standardize the data. The differentially expressed (DE) genes are listed in **Supplement Figure 2.**

Subsequently, the limma R package (*24*) was engaged to pinpoint differentially expressed genes (DEGs) within the R 4.4.1 (*25*) statistical environment.. The data were subjected to a Log2 transformation, and the stringency for identifying DEGs was set at an FDR-adjusted p-value of 0.15 or lower.

For GO functional enrichment analysis, we performed Gene Ontology (GO) enrichment analysis using the clusterProfiler package in R (*26–29*). Differentially expressed genes (DEGs) were identified based on the thresholds of log₂ fold change (log_2_FC) and adjusted p-value: genes with log_2_FC > 1 and adjusted p-value < 0.05 were considered upregulated, while genes with log_2_FC < -1 and adjusted p-value < 0.05 were considered downregulated. The org.Sc.sgd.db annotation package (specific for *Saccharomyces cerevisiae*) was installed via BiocManager) (*30*) and loaded into the R environment. GO enrichment was conducted with the enrichGO() function using the “Biological Process” (BP) ontology, with p.adjust < 0.05 as the significance threshold and the Benjamini–Hochberg method for multiple testing correction. To visualize the top enriched GO terms, we generated bubble plots using the top 20 significant GO terms based on adjusted p-values. Full list of GO could be found in **Supplement Figure 3.**

In parallel, we procured KEGG annotations from the Applied Protein Technology company and deployed the enrichKEGGfunction (*31*) to perform pathway enrichment analysis. From this analysis, we curated the top 20 KEGG pathways based on the adjusted P value, which were subsequently depicted using a dotplot for illustrative purposes. This visual representation facilitated a clear and concise interpretation of the pathways most significantly impacted by the observed gene expression changes. Illustrations are made by ImageGP2 (*32*)

### Metabolomics data analysis

Raw data were converted to .mzML format using ProteoWizard, followed by peak alignment, retention time correction, and peak area extraction using the XCMS programme. The metabolite structure identification was performed by searching the laboratory’s own database using exact mass number matching (<25 ppm) and secondary spectrum matching.

For the data obtained by XCMS extraction, ion peaks with >50% missing values within the group were removed. The software SIMCA-P 14.1 (Umetrics, Umea, Sweden) was applied for pattern recognition, and the data were pre-processed by Pareto-scaling and then subjected to multidimensional statistical analyses, including unsupervised Principal Component Analysis (PCA) analysis, supervised Partial least Squares discrimination analysis (Partial least squares discrimination analysis, PLS-DA) and Orthogonal partial least squares discrimination analysis (OPLS-DA). Unidimensional statistical analyses included Student’s t-test and multiples of variance analysis, and R software plotted volcano plots. According to the Variable Importance for the Projection (VIP) obtained from the OPLS-DA model to measure the intensity of the influence of the expression pattern of each metabolite on the classification discrimination of the samples in each group and the ability to explain, to search for biologically significant differential metabolites. In this experiment, VIP>1 was used as the screening criterion to preliminarily screen out the differentials among the groups. Univariate statistical analysis was further used to verify whether the differential metabolites were significant.

In order to evaluate the plausibility of candidate metabolites, and to more visualize the relationship between samples and the metabolite expression patterns, we utilized qualitatively significant differential metabolite expression to perform hierarchical clustering on each group of samples with R.

We used the KEGG “Search & Color Pathway” web tool (https://www.genome.jp/kegg/mapper.html) to identify the key changes as follows. First, metabolite identifiers were converted to KEGG compound IDs (e.g. C00049 for D-glucose), where positive values indicate up-regulation (red) and negative values down-regulation (green). High-resolution PNGs incorporated.

Metabolites with both multidimensional statistical analysis VIP>1 and univariate statistical analysis p<0.05 were selected as significantly different metabolites, while VIP>1 and 0.05<p<0.1 were considered as differential metabolites. Differential metabolites identified by the positive ion model are shown in Table S5, and those identified by the negative ion model are shown in Table S6.

## Results

### CSL increased the yeast fermentation quantity with physiology confirmation

We first examined the effect of yeast fermentation with additive of CSL in the medium. Fermentation was conducted with the concentration of CSL set at 2.69 mg/mL. After 18 hours added with CSL, the ethanol content in the yeast solution supplemented with CSL was measured at 44.6 ± 0.137 mg/mL, compared to 42.3 ± 0.037 mg/mL in the control group. Under the conditions of 2.69 mg/mL CSL concentration, the ethanol production by the yeast increased by 5.16% (**Figure 1** a).

**Figure 1.**
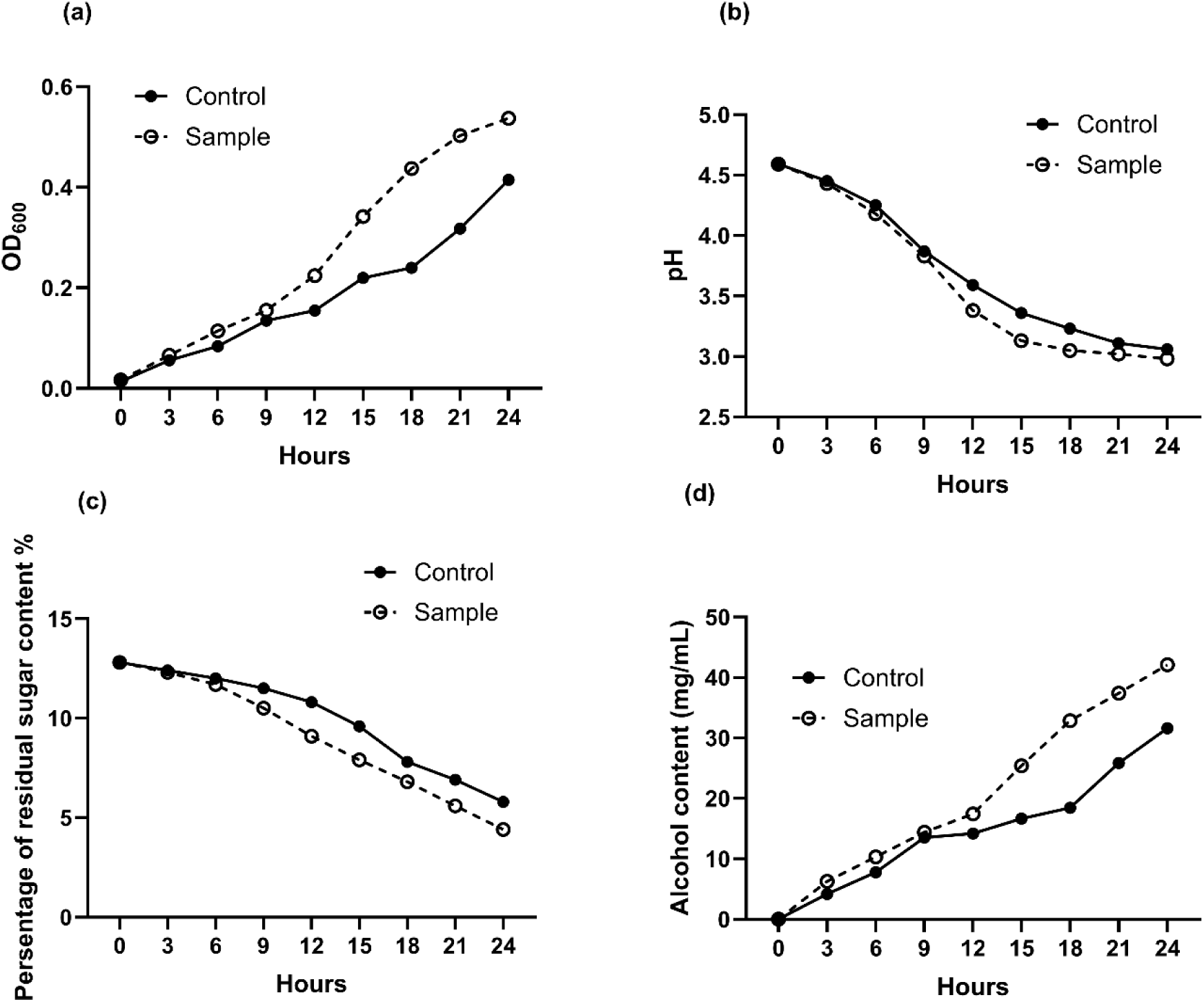
Effect of CSL on the growth of *Saccharomyces cerevisiae* during fermentation. (a) Effect of CSL on the proliferation of *Saccharomyces cerevisiae*. (b) Effect of CSL on pH value of fermentation broth. (c) Effect of CSL on residual sugar in fermentation broth. (d) Effect of CSL on ethanol concentration in fermentation broth.

During the fermentation of *Saccharomyces cerevisiae*, the pH, sugar level and ethanol yield all changed significantly (Figures 1 bcd). The pH of the fermentation broth decreased gradually with time, and the overall trend was the same between the control group and the CSL group. There was no significant difference in the change in the first 9 h, but the pH of the CSL group decreased more sharply after 9 h, which indicated that the CSL accelerated the decrease of pH in the fermentation process. Meanwhile, the consumption of glucose by brewer’s yeast gradually increased during the logarithmic growth period, and the glucose consumption of the CSL group was higher than that of the control group due to the higher number of yeast in the CSL group, which indicated that the CSL could increase the utilization of glucose by yeast, and thus affect the sugar content and taste of the wine. As the fermentation progressed, the ethanol production increased, and both groups showed an increasing trend. The ethanol content did not differ much in the first 9 h, but the ethanol production of the CSL group was significantly higher than that of the control group after 9 h, which indicated that the CSL could promote the synthesis of ethanol by brewer’s yeast to increase the alcoholic strength and improve the flavor of the wine.

### CSL enhances yeast fermentation by modulating membrane integrity, lipid biosynthesis, and energy metabolism

We carried out RNA-seq with *Saccharomyces cerevisiae* cultivated on blank medium with comparison of medium supplemented with lectin extracted from *Cyclina sinensis*.

*Saccharomyces cerevisiae* samples were quantified with qubit2.0 and further calibrated with qPCR and snap frozen with liquid nitrogen for sequencing. We carried out Differential Expression (DE) analysis with limma R package. From RNA-seq result, in total we found 871 upregulated genes and 959 downregulated genes (**Figure 2** b). We further used Goseq package (*33*) for Gene Ontology annotation with GO annotation from Applied Protein Technology company. The upregulation of several key pathways in both Molecular Function (MF) and Biological Process (BP) categories (as identified through GO analysis) is closely related to enhanced alcohol production in yeast. Details can be found in **Table S2-S3**.

**Figure 2.**
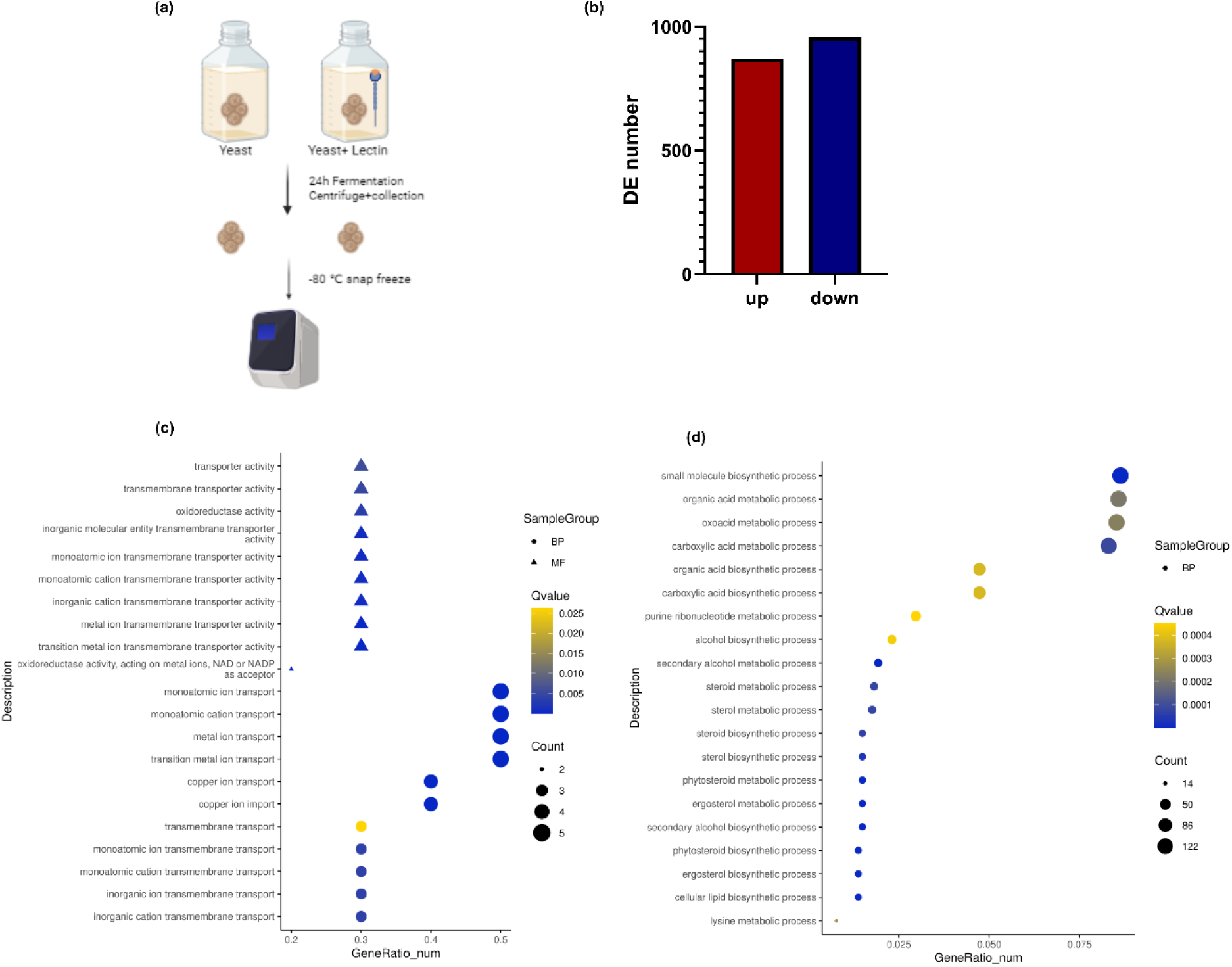
RNA sequencing Differential Expression (DE) and GO analysis results, highlighting multiple upregulated pathways related to alcohol production. (a) RNA-seq experimental design. The figure created by Biorender. (**b**) DE analysis summary: A summary of the differentially expressed (DE) genes, showcasing the total number of upregulated and downregulated genes. (**c**) Top 20 upregulated and downregulated (**d**) genes: A visualization of the top 20 significantly upregulated and downregulated genes, identified through DE analysis. The results of the GO analysis, focusing on three main categories: Molecular Function (MF), Biological Process (BP), and Cellular Component (CC). The bars represent the negative value of the adjusted P-values from the GO analysis. A detailed list of all GO analysis results can be found in Supplementary Data 1.

To elucidate the molecular changes induced by lectin treatment during yeast fermentation, Gene Ontology (GO) enrichment analysis was performed on the upregulated genes. The results revealed a significant enrichment in categories associated with metal ion transport, particularly copper ion import, iron ion transport, and zinc ion transport (Table S3). GO terms such as transition metal ion transmembrane transporter activity, monoatomic ion transport, and intracellular iron ion homeostasis were highly represented, suggesting an enhanced mobilization and regulation of essential metal ions under lectin treatment (Figure 2c) (*34,35*).

A second major functional theme involved oxidoreductase activity (GO:0016491) including oxidoreductase activity, acting on metal ions, acting on CH-OH group of donors, and NAD(P)H-dependent processes, is directly associated with enzymatic reaction in glycolysis.This enzymatic activity is pivotal for redox reactions within the glycolytic pathway, wherein glucose is catabolized into pyruvate, concurrently generating ATP and reducing equivalents that are subsequently utilized in ethanol biosynthesis. The upregulation of this function enhances the efficiency of glycolysis, maintaining redox balance, facilitating efficient conversion of carbon sources into ethanol, facilitating an augmented yield of ethanol (*36*). These oxidoreductases are critical for highlighting a potential mechanism glycolysis where lectin promotes alcohol biosynthesis (*37–39*) .In addition, GO terms linked to the Ehrlich pathway were significantly enriched, such as amino acid catabolic process via Ehrlich pathway, alcohol biosynthetic process, and alcohol metabolic process. These results indicate that amino acid degradation to fusel alcohols may be upregulated, further supporting enhanced ethanol and by-product formation (*40,41*).

GO enrichment analysis revealed that multiple core metabolic and biosynthetic processes were significantly downregulated in the lectin-treated group (Figure 2d, Supplement Table S3). The lipid biosynthetic process (GO:0008610) and the synthesis of very long-chain fatty acids (GO:0000038) are essential for constructing and maintaining the structural integrity of cell membranes. Lipids not only provide a matrix for embedding proteins and other molecules but also play a crucial role in modulating the physical properties of the membrane, including its thickness, curvature, and packing (*14,42*). Genes involved in fungal-type cell wall (GO:0009277) and cell wall organization or biogenesis (GO:0071554) were also downregulated, confirming possible interference with cell wall integrity (*14,43*). The cell membrane not only serves as a critical barrier regulating the influx and efflux of molecules but also plays a vital role in signal transduction and cellular homeostasis. The synthesis of cell membrane components involves several key biosynthetic processes, primarily focused on the production of ergosterol, other sterols, and various lipids (*44*). Ergosterol, analogous to cholesterol in animal cells, is a fundamental component of the fungal cell membrane that contributes significantly to the membrane’s fluidity and permeability. The ergosterol biosynthetic process (GO:0006696) and steroid biosynthetic process (GO:0006694) are central to maintaining cell membrane integrity in yeast (*45,46*). The downregulation of these pathways the yeast cells were suffering from ethanol toxicity with CSL, confirming the production promotion of fermentation with CSL.

Furthermore, energy-related pathways such as aerobic respiration (GO:0009060) and generation of precursor metabolites and energy (GO:0006091) were also inhibited, indicating a potential disruption of mitochondrial function and energy metabolism. Pathways includes small molecule biosynthetic process (GO:0044283), secondary alcohol metabolic process (GO:1902652), secondary alcohol biosynthetic process (GO:1902653), ergosterol biosynthetic process (GO:0006696), as well as amino acid metabolic process (GO:0006520), nucleotide metabolic process (GO:0009117), and organic acid metabolic process (GO:0006082). The downregulation of these pathways suggests a global suppression of anabolic activity under lectin treatment.

To identify significantly enriched pathways, we performed KEGG pathway en-richment analysis on both upregulated and downregulated DEGs. DEGs were mapped to their corresponding KEGG pathways using pathway annotations obtained from sequencing company. We then calculated the number of DEGs associated with each pathway and compared these counts to the total number of genes in each pathway using a hypergeometric test. P-values were adjusted for multiple testing using the Benjamini-Hochberg method to control for the false discovery rate (FDR), with adjusted P-values (FDR < 0.05) considered statistically significant. Separate analyses were con-ducted for upregulated and downregulated DEGs to identify distinct pathways affected in each case (**Figure 3**).

**Figure 3.**
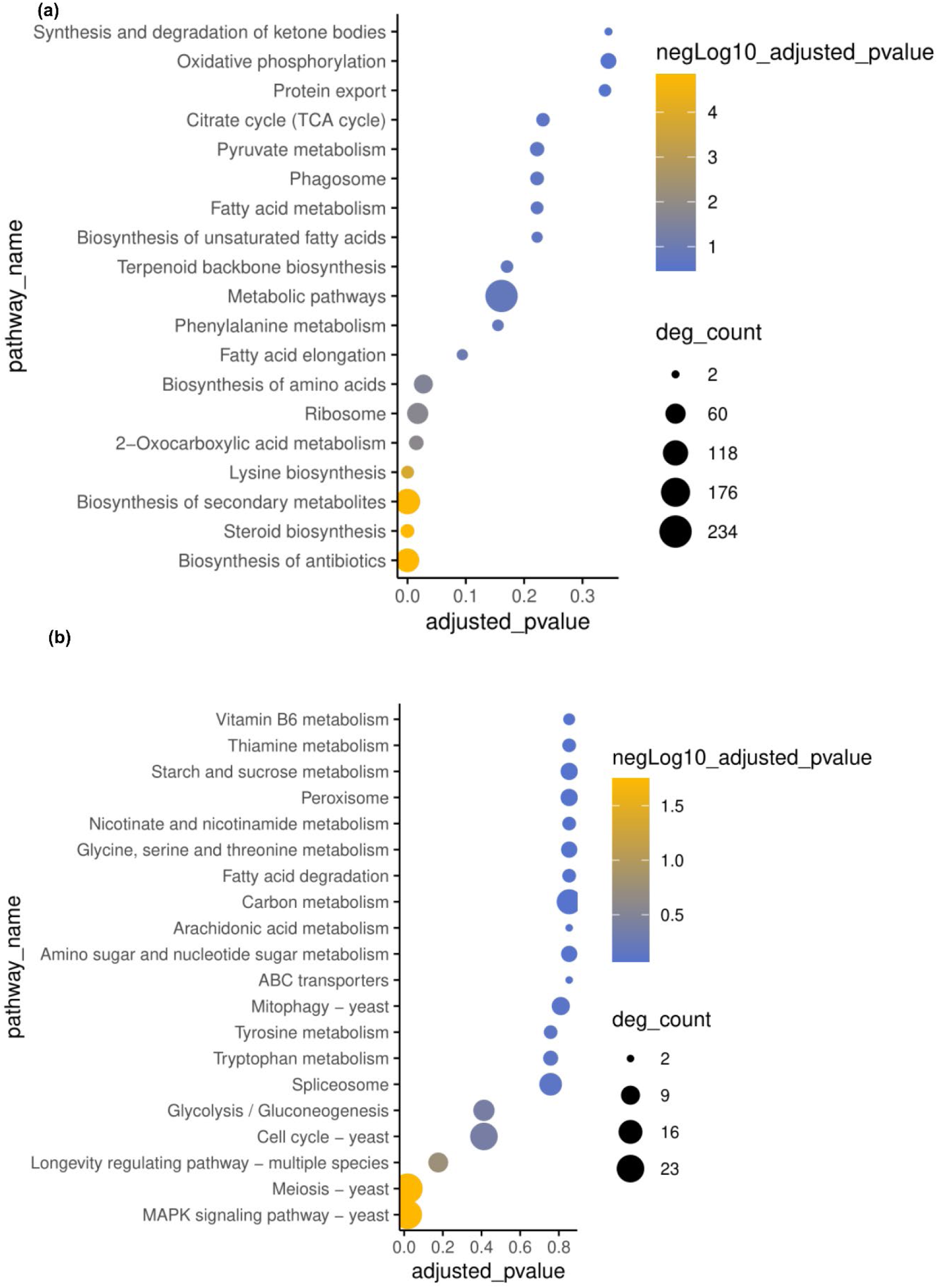
KEGG analysis of transcriptome data. (a) Upregulated DEGs with KEGG pathway enrichment, and (b) Downregulated DEGs with KEGG pathway enrichment. The dot size represents the number of differentially expressed genes (DEGs) associated with each pathway. The color scale indicates the negative log10 of the adjusted P-value, with yellow representing significant pathways where the adjusted P-value is less than 0.05.

Steroid biosynthesis (adjusted P-value: 1.47E-05) plays a role in maintaining the integrity of cell membranes, enhances membrane integrity under stress, particularly from high ethanol concentrations, suggesting an adaptive mechanism to stabilize cell membranes with sterol molecules like ergosterol (Hoshino and Gaucher, 2021). In pathway Lysine Biosynthesis (adjusted P-value: 0.000157775), lysine is crucial for protein synthesis and cellular growth, lysine biosynthesis may support the increased demand for protein synthesis as yeast cells proliferate under fermentation conditions (Isogai et al., 2021; Van Den Brink et al., 2008).

Additionally, lysine is a precursor in various metabolic pathways that contribute to yeast’s ability to thrive in nutrient-limited environments like those encountered during fermentation.

Biosynthesis of secondary metabolites (adjusted P-value: 1.59E-05) indicates a shift in metabolism towards the production of ethanol and related by-products, crucial for fermentation efficiency. Biosynthesis of Antibiotics (adjusted P-value: 1.47E-05) likely reflects a defense strategy against microbial competitors, enhancing yeast dominance during alcohol fermentation (Awan et al., 2017). 2-Oxocarboxylic acid metabolism (adjusted P-value: 0.015075274) and Biosynthesis of amino acids (adjusted P-value: 0.027223502) are upregulated to optimize central carbon metabolism and amino acid production, respectively, ensuring sufficient energy and biomass for sustained fermentation. Ribosome Biogenesis (adjusted P-value: 0.01747016) underscores the increased need for protein synthesis as yeast undergoes rapid growth and metabolic activity essential for alcohol production.

MAPK signaling pathway (adjusted P-value: 0.017920251) and Meiosis (adjusted P-value: 0.017920251) are downregulated, suggesting a reduced need for stress response and sexual reproduction under fermentation conditions. This shift favors rapid asexual reproduction to maximize population growth and alcohol output. Overall, these pathway changes reflect yeast’s strategic metabolic adjustments to optimize alcohol production while maintaining cellular integrity and growth under the demanding conditions of fermentation.

### Metabolic profiling of CSL-activated yeast reveals distinct metabolic signatures

UHPLC-Q-TOF MS was used for the separation and data acquisition of CSL-activated yeast (T) and control yeast (C) samples. Typical total ion chromatogram (TIC) profiles of CSL-activated yeast and control yeast analyzed by UHPLC-Q-TOF MS are shown in **Supplementary Figure 1**. Comparison of spectral overlap was performed for the total ion chromatograms, and as seen in the figure, the response intensities and retention times of the peaks of each chromatogram for each group of samples were basically overlapped, indicating that the parallelism between the two groups of samples was good.

Multivariate statistical analyses, including PCA, PLS-DA, and OPLS-DA, were performed to explore metabolic differences between CSL-reacting yeast (T) and control yeast (C) groups. PCA score plots (**Supplementary Figure 2**) revealed a clear separation between the groups in both positive and negative ion modes, confirming distinct metabolic profiles. The robustness of the PCA model was supported by the parameters obtained after 7-fold cross-validation (Table S7). PLS-DA further demonstrated the relationship between metabolite expression and sample categorization, with high R²Y and Q² values (0.989 and 0.972 for the positive ion model; 0.993 and 0.927 for the negative ion model), indicating strong model reliability (**Supplementary Figure 3**, **Table S8**). Additionally, OPLS-DA (**Supplementary Figure 4**) effectively discriminated between Group T and Group C, with R²Y and Q² values of 0.989 and 0.965 for the positive ion model, confirming the stability and predictive power of the model (**Table S9**). These results highlight the significant metabolic alterations induced by CSL in yeast. As illustrated, no overfitting occurred in the OPLS-DA model for either mode.

### CSL enhances ethanol fermentation in *Saccharomyces cerevisiae* by modulating glycolysis, TCA Cycle, and redox metabolism

To investigate the impact of CSL on yeast fermentation metabolism, we performed negative-ion-mode metabolomics and hierarchical clustering, revealing that CSL treatment depleted upstream substrates (e.g., sugar phosphates and TCA intermediates) while elevating downstream organic acids, NAD(P)H, and Ehrlich-pathway products; notably, CSL significantly increased key glycolysis and TCA steps (eg. Citrate, L-Malic acid, Succinate, L-Glutamate, L-Aspartate). It is hypothesized that it is not primarily involved in energy production, but more likely reflects reprogramming of metabolic fluxes that may serve functions such as carbon skeleton replenishment, biosynthesis, redox homeostasis maintenance, or cellular stress response, thereby enhancing flavorful organic acid production with high value for brewing applications (**Figure 4, 5**) (**Table S4,S5, S6**) (*47–50*).

**Figure 4.**
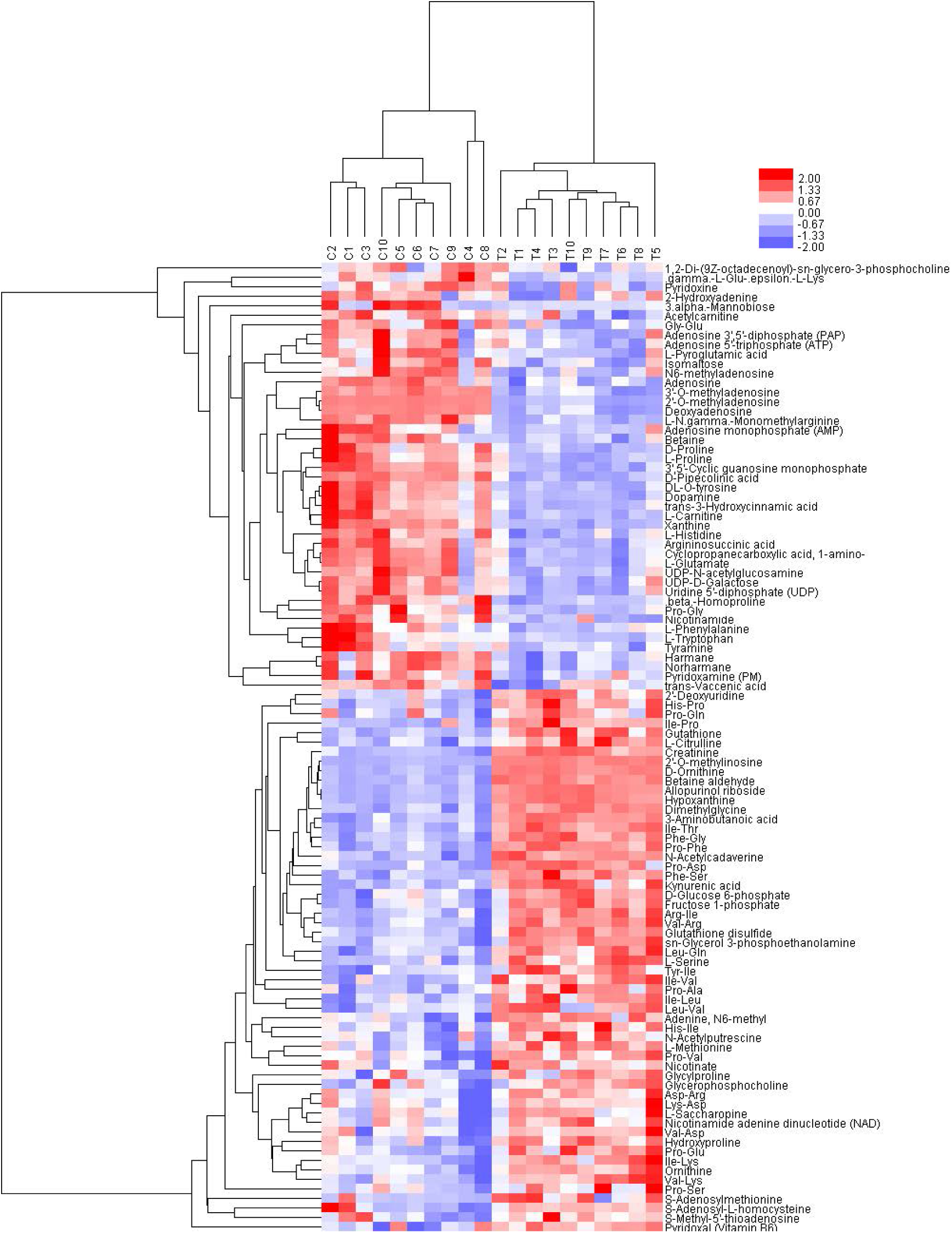
Hierarchical clustering analysis of significantly different metabolites in positive ion mode. This heatmap represents the hierarchical clustering analysis of significantly different metabolites between the control group (C, yeast without lectin treatment) and the treatment group (T, yeast treated with lectin) under positive ion mode. The red color indicates metabolites with higher relative abundance, while the blue color represents those with lower relative abundance.

*Saccharomyces cerevisiae* absorbs glucose from the medium for intracellular, glucose is converted to glucose 6-phosphate by the glycolytic pathway, and glucose 6-phosphate is then cleaved into dihydroxyacetone phosphate and glyceraldehyde 3-phosphate through a series of transformations and eventually converted into pyruvate, which is converted to ethanol by alcoholic fermentation. Glucose is an important carbon source for yeast cell fermentation and a signaling molecule in the early stages of fermentation (*51*). The samples of group T and C were analyzed and 98 were obtained as significantly different metabolites (p<0.05) by positive ion mode, of which 55 were up-regulated and 43 were down-regulated. Seventy-nine were obtained as significantly different metabolites (p<0.05) by negative ion mode, of which 64 were up-regulated and 15 down-regulated. Among these metabolites, the treatment group (T) exhibited significantly increased levels of alpha-D-glucose, D-glucose 6-phosphate, dihydroxyacetone phosphate, and citrate. These results suggest a shift in carbon flux, with potential enhancement of glycolytic intermediates accumulation under the treatment condition. (Figure 4) (Table S5).

Metabolomics analysis showed that the addition of CSL significantly elevated the levels of key intermediate metabolites (e.g., cis-aconitic acid, succinic acid, and malic acid) in the glycolysis and tricarboxylic acid cycle (TCA) pathways of *Saccharomyces cerevisiae*, promoting the production of flavorful organic acids, which is of great value for application in the brewing industry.

Nicotinamide adenine dinucleotides (NAD^+^ and NADH) are unable to cross the mitochondrial membrane of *S. cerevisiae*, and their oxidative and reductive interconversions rely on the malate-aspartate shuttle system for “shuttling”(*52*). In terms of reducing power balance, the contents of NAD^+^, malate and aspartate in the metabolites of CSL-supplemented yeast cells were significantly increased compared with those of the control group, suggesting that CSL promotes the inter-conversion of NAD^+^ and NADH and accelerates the rate of oxidation-reduction reactions. In addition, the elevated intracellular glucose content in the CSL group induced the Crabtree effect, which reduced the activity of the TCA pathway and enhanced the EMP pathway, leading to a large accumulation of ethanol. During alcoholic fermentation in *S. cerevisiae*, although there was no net surplus of total NADH, the high concentration of glucose induced the EMP pathway to produce 2 molecules of NADH in synchronization with the metabolism of 1 molecule of glucose to 2 molecules of pyruvate, thus generating an additional reducing power.

In addition, dihydroxyacetone phosphate accepts hydrogen on NADH to produce 3- phosphoglycerol, with concomitant release of NAD^+^, resulting in elevated levels of 3- phosphoglycerol and NAD^+^ in metabolites. This phenomenon is further supported by the significantly higher levels of lactate and 3-phosphoglycerol in the CSL group compared to the control group. When in negative ion mode, there was more DL- lactate expression in the CSL group, in positive ion mode, lactate was not detected (or not significant), this is possibly because the other bacterium, not yeast, or a false positive test.

Glutathione (GSH) consists of glutamic acid, cysteine and glycine linked by an amide bond and has strong antioxidant properties, scavenging free radicals in the body and protecting active sulfhydryl proteins from oxidative damage (*53*). Its oxidized form is glutathione disulfide (GSSG), which consists of two molecules of GSH linked by dehydrogenation to form a disulfide bond. In terms of nitrogen metabolism, during ethanol fermentation in *Saccharomyces cerevisiae*, GSH serves as a concomitant product, and its content increases with ethanol production (*54,55*). In terms of reducing power balance, the contents of 3-phosphoglycerol was significantly increased in the CSL group, indicating that the excess NADH produced during glycolysis was mainly consumed through the dihydroxyacetone phosphate and pyruvate metabolism pathways to generate 3-phosphoglycerol, thus maintaining the intracellular redox homeostasis. In addition, CSL treatment significantly elevated the content of glutathione (GSH) and its oxidized form (GSSG), and enhanced the antioxidant capacity of the cells, which might be related to the oxidative stress caused by ethanol accumulation.

In nitrogen metabolism, ornithine, citrulline and arginine are intermediate metabolites in the urea cycle. Under the action of ATP, CO_2_ and NH_3_ were catalysed by carbamoyl phosphate synthase to generate carbamoyl phosphate, which entered the urea cycle and released fumaric acid and urea with the participation of aspartic acid (*53*).The contents of ornithine, citrulline, and arginine were significantly increased in the CSL group, suggesting that the CSL effectively reduced ammonia toxicity and optimized the efficiency of nitrogen metabolism by promoting the urea cycle. Meanwhile, these metabolic changes not only improved the ethanol fermentation efficiency, but also promoted the accumulation of flavor substances (e.g., cis-uronic acid, succinic acid, malic acid), which is important for winemaking flavor regulation.

In negative ion mode, Cluster 1 (Down in CSL; high in control) metabolites include many primary carbon sources: glucose-6-phosphate, fructose-1,6-bisphosphate, citrate, malate, several amino acids (serine, threonine), and nucleotides (adenosine, guanosine) (Figure 5)(Table S6).

**Figure 5.**
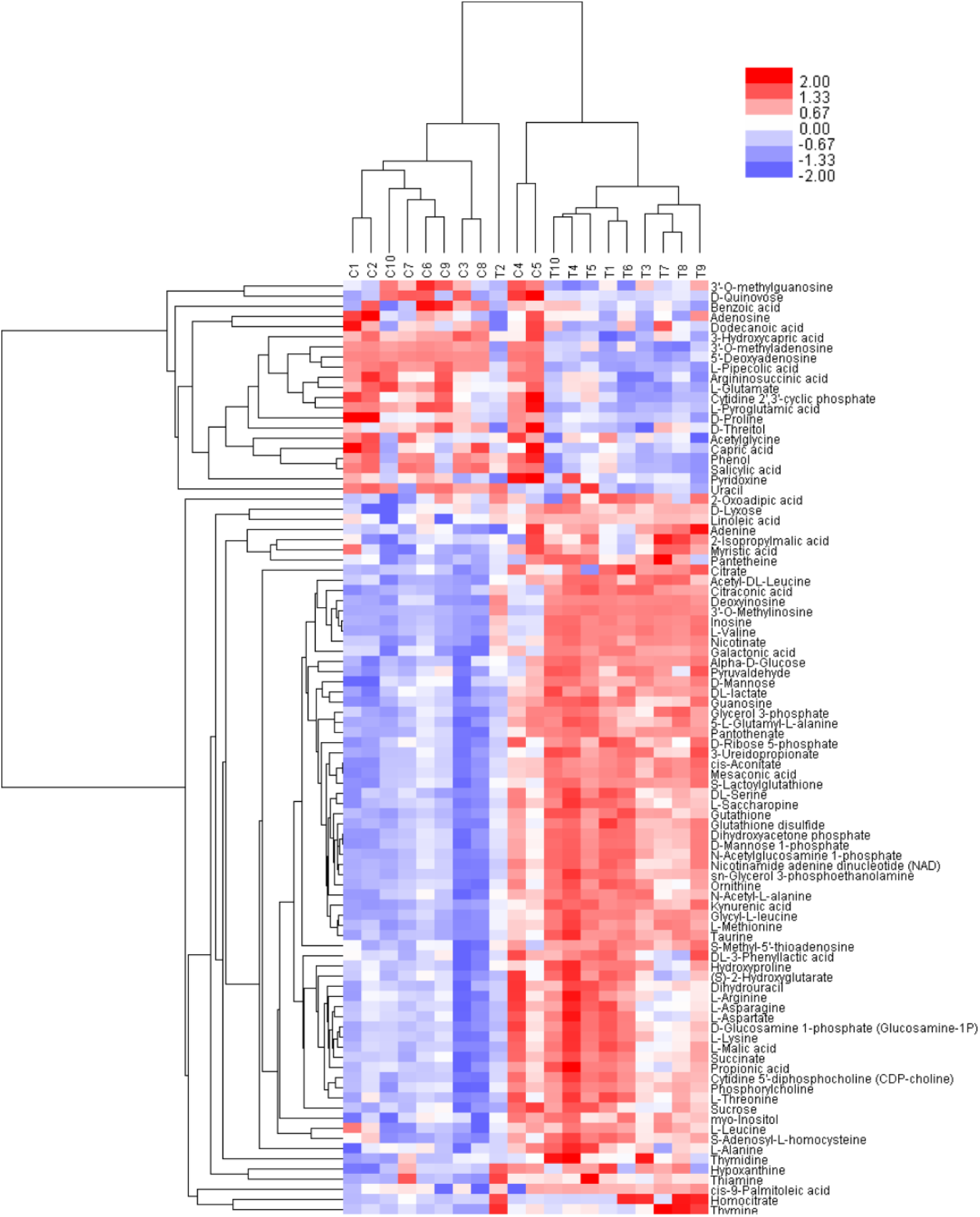
Hierarchical clustering analysis of significantly different metabolites in negative ion mode. This heatmap represents the hierarchical clustering analysis of significantly different metabolites between the control group (C, yeast without lectin treatment) and the treatment group (T, yeast treated with lectin) under negative ion mode. The red color indicates metabolites with higher relative abundance, while the blue color represents those with lower relative abundance.

Their depletion under CSL suggests that lectin treatment accelerates their consumption, funneling them into downstream pathways (e.g. glycolysis, TCA cycle, amino acid catabolism).

Cluster 2 (Up in CSL; low in control) is enriched for downstream catabolites and redox cofactors, including pyruvate, NAD/NADH, various organic acids (2-oxoglutarate, succinate), Ehrlich- pathway fusel alcohol precursors, and lipid breakdown products, which further confirmed our finding from GO analysis. Their accumulation in CSL treated group indicates an enhancement of glycolytic flux and activation of the Ehrlich pathway, together reveals that CSL treatment rewires yeast central metabolism. CSL both depletes upstream substrates and boosts the pathways that convert them into ethanol and by-products.

In order to visualization DE changes in regulation pathways, we illustrated the key pathways in Figure 6. In glycolysis/gluconeogenesis pathway (KEGG map00010), metabolites including α-D-glucose and glycerone-P (DHAP)—were significantly upregulated upon CSL treatment (Figure 6 a). α-D-glucose indicates intracellular accumulation of free glucose or its phosphorylated precursors, likely due to enhanced external sugar uptake or restricted downstream flux (*56*).

**Figure 6.**
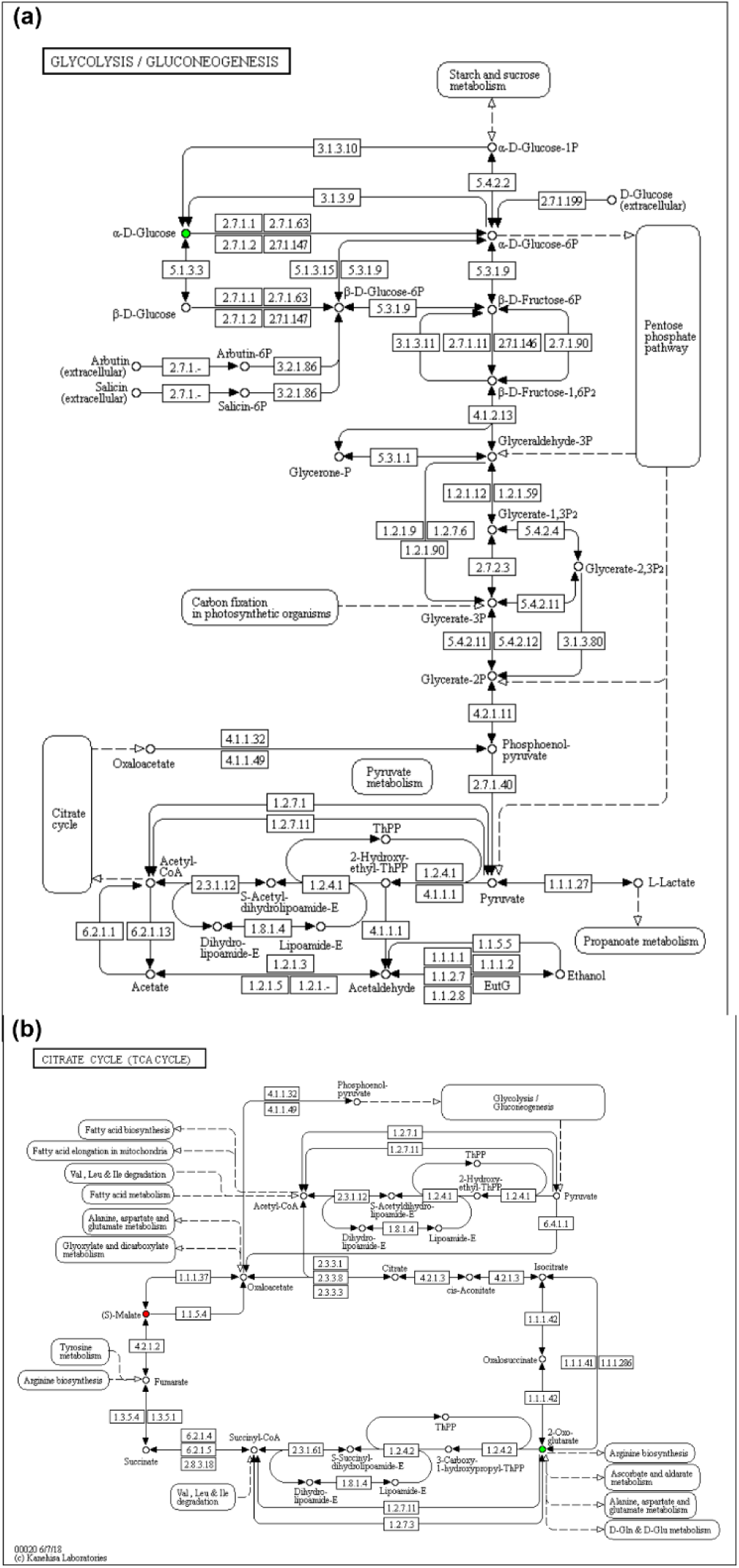
KEGG pathway maps of central carbon metabolism in *Saccharomyces cerevisiae* s upon CSL treatment. (a) Glycolysis/gluconeogenesis pathway (map00010). (b) Tricarboxylic acid (TCA) cycle pathway (map00020). Nodes represent metabolites, colored by log_2_-fold change (green= down-regulated; white = no change; red = up-regulated).

Glycerone-P (DHAP) suggests a bottleneck at the GAPDH step (EC 1.2.1.12) after fructose-1,6-bisphosphate cleavage (EC 4.1.2.13), leading to significant pooling of triose fragments.These shifts collectively suggest that GAPDH/NADH redox balance becomes a critical regulatory node, driving both fermentative efficiency and byproduct diversification.

Similarly, the TCA-cycle map (map00020) dovetails perfectly with the glycolysis/gluconeogenesis shifts we saw in map00010.An increased flux from acetyl-CoA into the TCA cycle’s first steps (catalyzed by citrate synthase and aconitase), consistent with a higher delivery of pyruvate-derived acetyl-CoA from glycolysis. Citrate and cis-aconitate (GO:0006101 & GO:0006102) indicates they are significantly elevated under CSL treatment. TCA intermediates—Succinate and (S)-malate (GO:0006103 & GO:0006104) also show upregulated, meaning they accumulate in the CSL-treated cells (*48,57*).

Elevation of these TCA intermediates, together with the glycolytic shifts, points to an overall upregulation of central carbon catabolism. Combining proteomics and metabolomics data, CSL significantly enhanced the fermentation performance of *S. cerevisiae* through multi-level metabolic regulation, including glycolysis, redox balance, nitrogen metabolism and antioxidant defense. These findings provide theoretical support for the application of CSL in the brewing industry and lay the foundation for further optimization of the fermentation process.

## Discussion

In this study, we systematically analyzed the effects of CSL on the metabolism and transcriptome of *Saccharomyces cerevisiae* by combining metabolomics and transcriptomics techniques. The results indicated that CSL significantly enhanced yeast fermentation performance by enhancing the glycolytic pathway, inhibiting the tricarboxylic acid cycle (TCA) and promoting ethanol accumulation (*39,58,59*). In addition, CSL enhanced yeast tolerance during fermentation by regulating redox homeostasis and antioxidant defense mechanisms.

These findings provide new insights into the role of CSL in the regulation of yeast metabolism and lay a theoretical foundation for its application in industrial fermentation.

Metabolomics data showed that α-D-glucose and glucose-6-phosphate contents were significantly up-regulated in the CSL-treated group, indicating that the glycolysis pathway was significantly enhanced. Meanwhile, key metabolites in the TCA cycle (e.g. malate, aspartate) were significantly accumulated, suggesting that the TCA cycle activity was inhibited. The transcript levels of ethanol fermentation-related enzymes were also significantly up-regulated, further confirming the promotional effect of CSL on ethanol synthesis. This phenomenon may be related to the fact that CSL affects the shunting of the sugar metabolic pathway, yeast prefers to generate ethanol via the glycolytic pathway rather than complete oxidation.

Metabolomics analysis showed a significant increase in NAD^+^, malate and aspartate in the CSL-treated group, suggesting a decrease in TCA cycling activity. Transcriptomic data further indicated that the expression of key genes of the TCA cycle was significantly down-regulated.

Possible mechanisms include CSL-mediated modulation of mitochondrial function that reduces cellular dependence on oxidative phosphorylation, thereby reprogramming energy metabolism to a more glycolysis-dependent rapid ATP generation pathway. This metabolic reprogramming not only increased ethanol production, but also potentially reduced the cellular requirement for TCA cycle-dependent growth pathways, thereby optimizing the metabolic efficiency of yeast under fermentation conditions (*60–62*). Ethanol fermentation is usually accompanied by reactive oxygen species production to alleviate oxidative stress in cells, which is both presented from RNA-sequencing and metabolism analysis(*60,63*). These findings suggest that CSL not only regulates sugar metabolism, but also helps yeast adapt to high ethanol concentrations by enhancing the antioxidant defense system.

As a GalNAc/Man binding lectin, CSL may interact with yeast cell surface glycoproteins to trigger metabolic reprogramming within the cell. This interaction may regulate sugar metabolism and mitochondrial function. In addition, CSL may help yeast resist ethanol toxicity by regulating cell membrane integrity. Future studies could further explore the targets of CSL’s direct action and its specific mechanism in metabolic regulation through gene knockdown or protein interactions analysis.

This study reveals the mechanism by which CSL significantly enhances yeast fermentation performance through multilevel metabolic regulation, providing important clues for the application of lectins in microbial metabolic regulation. These findings not only help to understand the role of CSL in yeast metabolism, but also provide theoretical support for the optimization of ethanol production in industrial fermentation. Future studies can further explore the potential application of CSL in other industrial microorganisms and combine proteomics and gene editing technologies to deeply resolve CSL-mediated signaling pathways and their broader roles in metabolic regulation. In addition, the role of CSL in regulating the accumulation of flavor substances (e.g., succinic acid, malic acid) deserves further investigation to optimize flavor regulation in winemaking processes.

Our study shows that CSL significantly affects the metabolism of *Saccharomyces cerevisiae* by enhancing glycolysis, inhibiting the TCA cycle and promoting ethanol accumulation. These metabolic changes were accompanied by an enhanced antioxidant stress response, which may help yeast adapt to the CSL-induced high ethanol environment. Future studies may further explore the CSL-mediated signaling pathways and their broader roles in metabolic regulation, providing more theoretical basis and technical support for the application of lectins in industrial fermentation.

## Conclusion

Our study demonstrates that *Cyclina sinensis* lectin (CSL) enhances ethanol production in *Saccharomyces cerevisiae* by modulating key metabolic pathways, including glycolysis and the TCA cycle. These findings provide a theoretical basis for the potential application of CSL as a natural fermentation enhancer in the brewing industry.

## Data availability

The raw RNA-sequencing data are available in the Gene Expression Omnibus (GEO) database under accession number GSE295666. The metabolomics data have been deposited in the MetaboLights repository under study identifier MTBLSREQ20250508210379.

## Supporting Information

The following files are available free of charge.

Table S1. Raw exon counts of RNA-seq analysis samples (.xlsx).

Table S2. Summary of differentially expressed genes (DEGs) (.xlsx).

Table S3. Gene ontology (GO) enrichment analysis (.xlsx).

Table S4. Characteristics of 117 proteins that were differentially expressed in S. cerevisiae following treatment with CSL (.xlsx).

Table S5. Differential metabolites identified in positive ions modes (.xlsx).

Table S6. Differential metabolites identified in negative ions modes (.xlsx). Table S7. Parameters of PCA (.xlsx).

Table S8. Evaluation parameters of PLS-DA in positive and negative ions modes (.xlsx).

Table S9. Evaluation parameters of OPLS-DA in positive and negative ions modes (.xlsx).

Figure S1. TIC chromatograms of S. cerevisiae under different conditions. (a) Positive ion mode for CSL-treated group (1T), (b) Positive ion mode for control group (2C), (c) Negative ion mode for CSL-treated group (3T), (d) Negative ion mode for control group (4C) (Tif).

Figure S2. Score plots of PCA in positive and negative ions modes. (a) Positive ion mode. (b) Negative ion mode (Tif).

Figure S3. Score plots of PLS-DA in positive and negative ions modes. (a) Positive ion mode. (b) Negative ion mode (Tif).

Figure S4. OPLS-DA analysis of S. cerevisiae metabolic profiles. (a) Score plot of OPLS-DA in positive ion mode. (b) Score plot of OPLS-DA in negative ion mode. (c) Permutation test of OPLS-DA in positive ion mode. (d) Permutation test of OPLS-DA in negative ion mode(Tif).

## Author Contributions

The manuscript was written through contributions of all authors. All authors have given approval to the final version of the manuscript. ‡These authors contributed equally.

## Funding Sources

This work was funded by the Natural Science Foundation of Inner Mongolia Autonomous Region (Grant no.2021MS03080) and the National Natural Science Foundation of China (Grant No. 31571916).

## ACKNOWLEDGMENT

We want to express our gratitude to Shanghai Applied Protein Technology Co., Ltd. for technical support in this work.

## ABBREVIATIONS

CSL: Cyclina sinensis lectin

## Notes

### Competing Interest Statement

The authors have declared no competing interest.

