## Supplementary figures and images for "Metabolic and transcriptomic insights into a GalNAc/Man-Specific lectin in yeast fermentation"

### Figure S1.tif

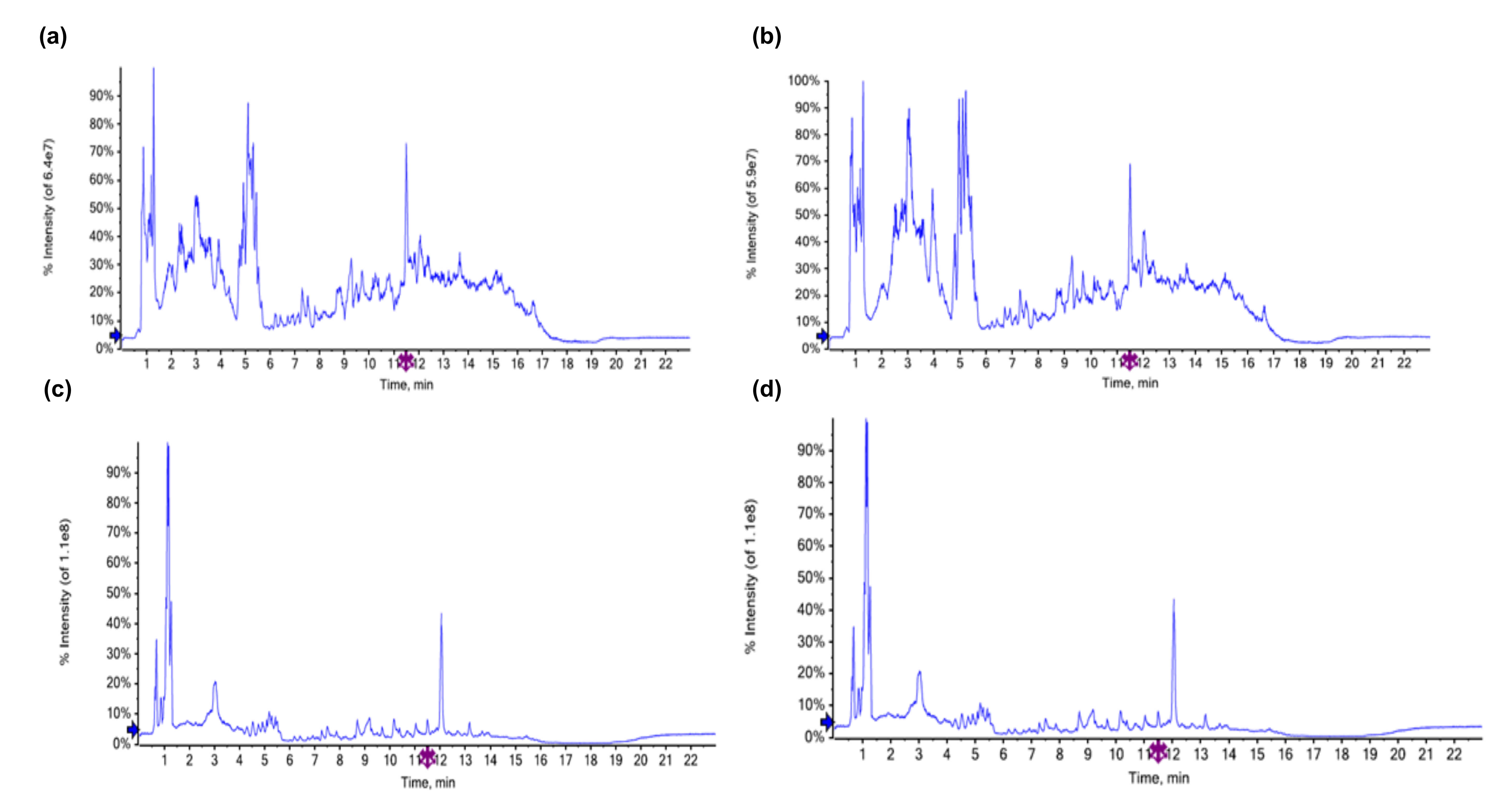

### Figure S2.tif

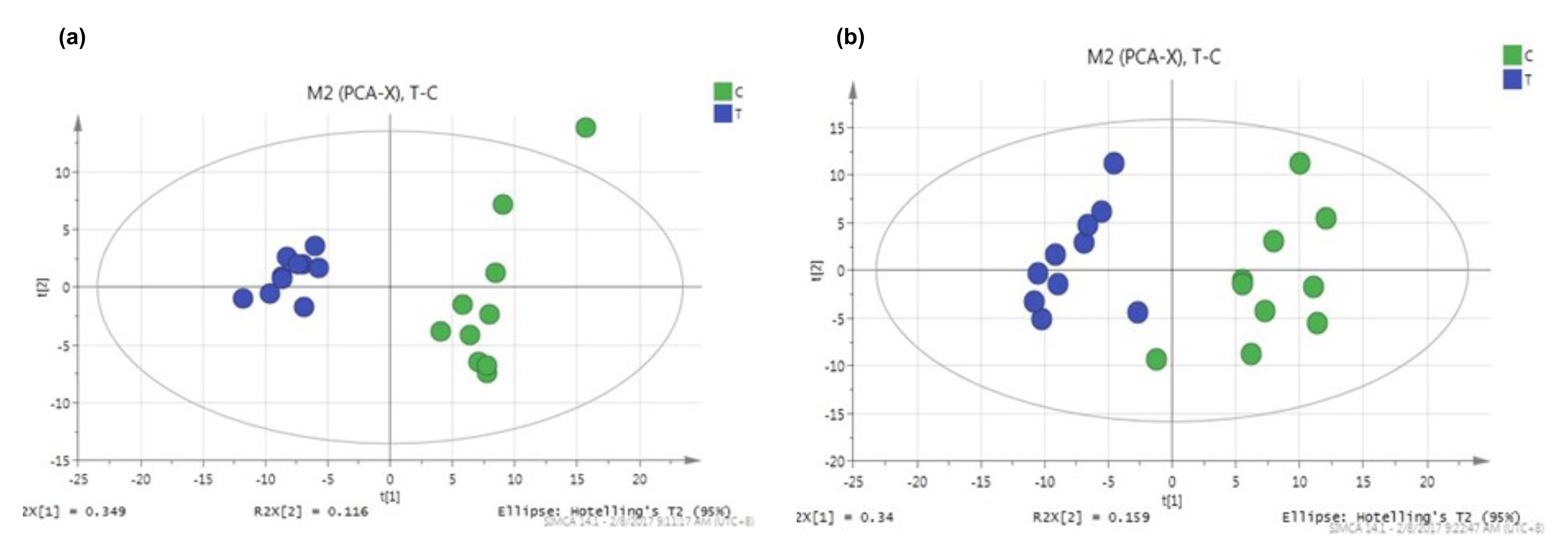

### Figure S3.tif

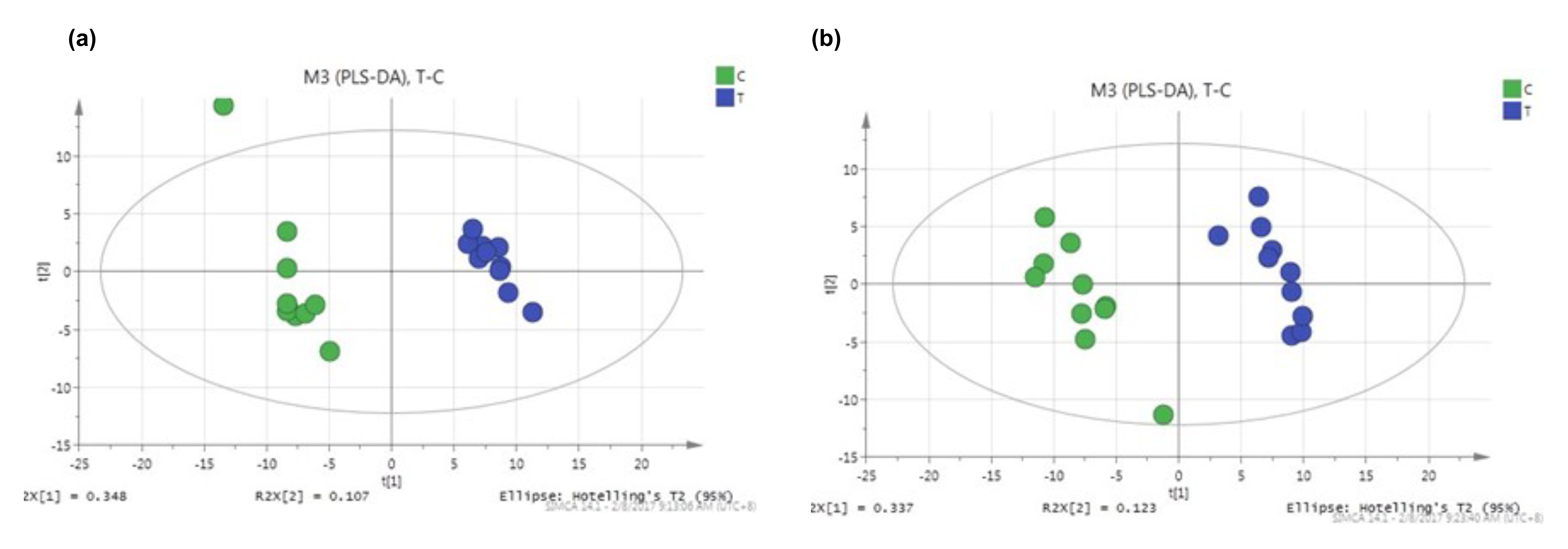

### Figure S4.tif

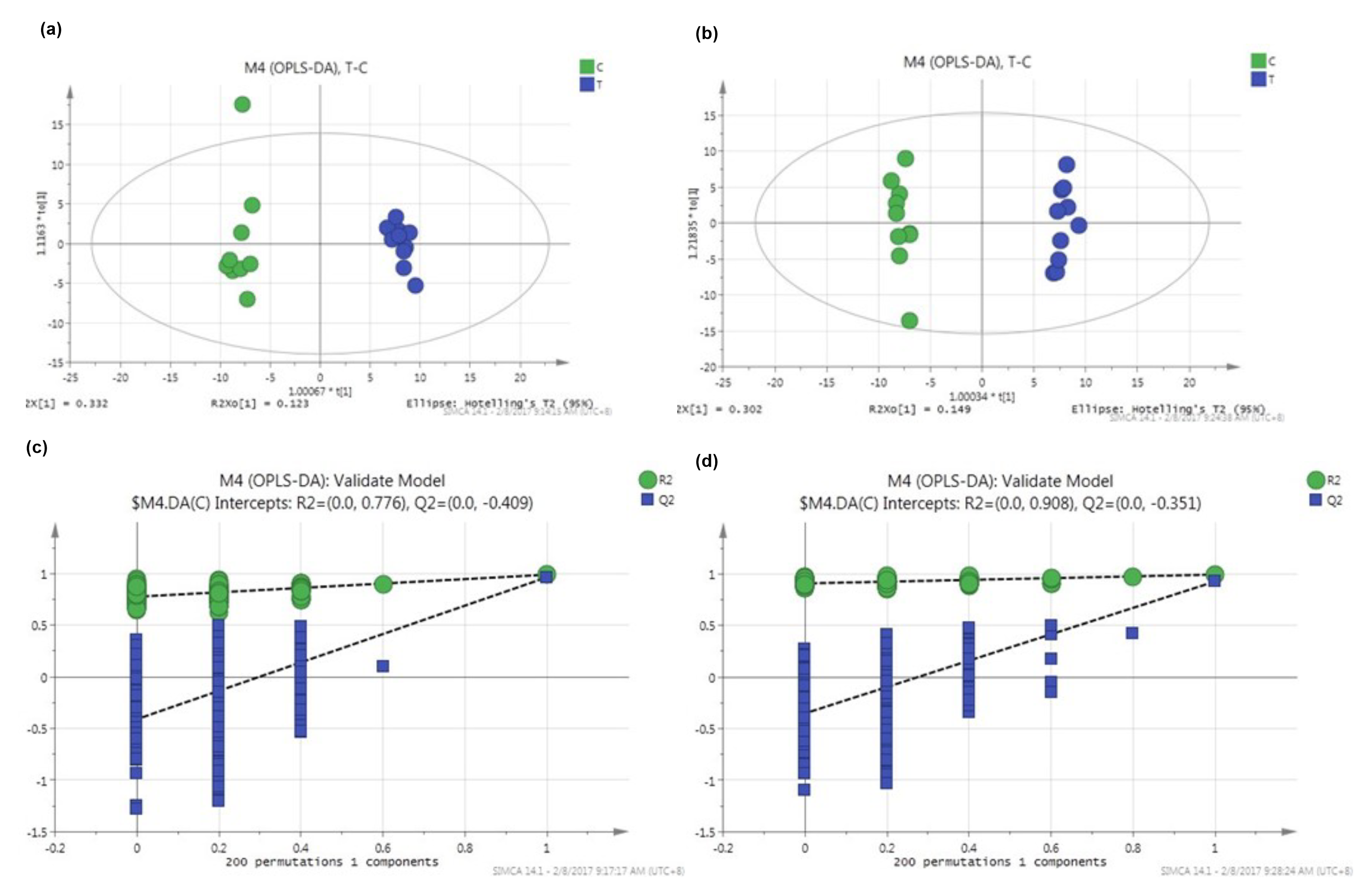
